# Shared enhancer gene regulatory networks between wound and oncogenic programs

**DOI:** 10.1101/2022.06.17.496596

**Authors:** Swann Floc’hlay, Ramya Balaji, Valerie Christiaens, Carmen Bravo González-Blas, Seppe De Winter, Gert Hulselmans, Maxime De Waegeneer, Xiaojiang Quan, Duygu Koldere, Mardelle Atkins, Georg Halder, Anne Classen, Stein Aerts

## Abstract

Wound response programs are often activated during neoplastic growth in tumors. In both wound repair and tumor growth, cells respond to acute stress and balance the activation of multiple programs including apoptosis, proliferation, and cell migration. Central to those responses are the activation of the JNK/MAPK and JAK/STAT signaling pathways. Yet, to what extent these signaling cascades interact at the *cis*-regulatory level, and how they orchestrate different regulatory and phenotypic responses is still unclear. Here, we aim to characterize the regulatory states that emerge and cooperate in the wound response, using the *Drosophila melanogaster* wing disc as a model system, and compare these with cancer cell states induced by *ras*^V12^*scrib*^-/-^ in the eye disc. We used single-cell multiome profiling to derive enhancer Gene Regulatory Networks (eGRNs) by integrating chromatin accessibility and gene expression signals. We identify a “proliferative” eGRN, active in the majority of wounded cells and controlled by AP-1 and STAT. In a smaller, but distinct population of wound cells, a “senescent” eGRN is activated and driven by C/EBP-like transcription factors (Irbp18, Xrp1, Slow border, and Vrille) and Scalloped. In tumor cells on the other hand, both eGRN signatures are simultaneously active within the same cell. Our single-cell multiome and eGRNs resource offers an in-depth characterisation of the senescence markers, together with a new perspective on the shared gene regulatory programs acting during wound response and oncogenesis.

## Introduction

The Drosophila wing imaginal disc (WID) is a classical model system to study developmental patterning and cell differentiation. This larval primordium is composed of two epithelial cell layers, the peripodial epithelium and the disc proper, along with muscle precursors (Figure 1.a). Despite its rather complex structure, the wing disc is extensively studied for its regeneration capacities. Damaged wing discs can trigger a set of wound-response mechanisms allowing for disc repair and the formation of normal wings (Smith-Bolton, 2016; Tripathi and Irvine, 2022). The molecular pathways identified as key drivers for this regenerative process include the regulation of cell apoptosis (JNK, JAK/STAT (Fortezza et al., 2016)), cell proliferation (EGFR, Wnt, Wingless, Scalloped (Blanco et al., 2010; Herrera et al., 2013; Smith-Bolton et al., 2009; Yu et al., 2015)), re-epithelialization (ERK, Grainy head (Mace et al., 2005)) and developmental timing (Insulin-like peptide 8 (Katsuyama et al., 2015)). Interestingly, the same regulatory pathways may lead to uncontrolled cell apoptosis or neoplastic growth when unrestricted (La Marca and Richardson, 2020; Pérez-Garijo et al., 2013; Pinal et al., 2019). Such tumor-like outcomes are observed in *ras*^V12^*scrib*^-/-^ transformed cells, where the loss of cell polarity triggers a cellular stress that cells cannot escape via apoptosis (Atkins et al., 2016; Brumby and Richardson, 2003; Cosolo et al., 2019; Davie et al., 2015; Fortezza et al., 2016; Igaki et al., 2006; Külshammer et al., 2015; Pagliarini and Xu, 2003; Pinal et al., 2019; Uhlirova and Bohmann, 2006). In light of these outcomes, it is still unclear how such antagonistic mechanisms (apoptosis and proliferation) interact in the vicinity of a wound to orchestrate tissue repair while controlling for overproliferation. To study this process at the gene regulatory level, we generated a multi-omic dataset, jointly measuring chromatin accessibility and gene expression changes at single-cell resolution, in wild-type and genetically ablated wing imaginal discs from 3rd instar larvae (Figure 1.a).

Leveraging the multidimensionality of our dataset, we constructed enhancer gene regulatory networks (eGRN), centered around Transcription Factors (TF) and comprising both gene and enhancer signatures (Janssens et al., 2022). We detect two classes of wound populations, respectively expressing markers of proliferation and cellular senescence. The senescent cells are driven by eGRNs belonging to the C/EBP bZip family (Irbp18, Slow border, Vrille (Blanco et al., 2020)), which we also found to be present in the tumor cells from the *ras*^V12^*scrib*^-/-^ model.

**Figure 1:**
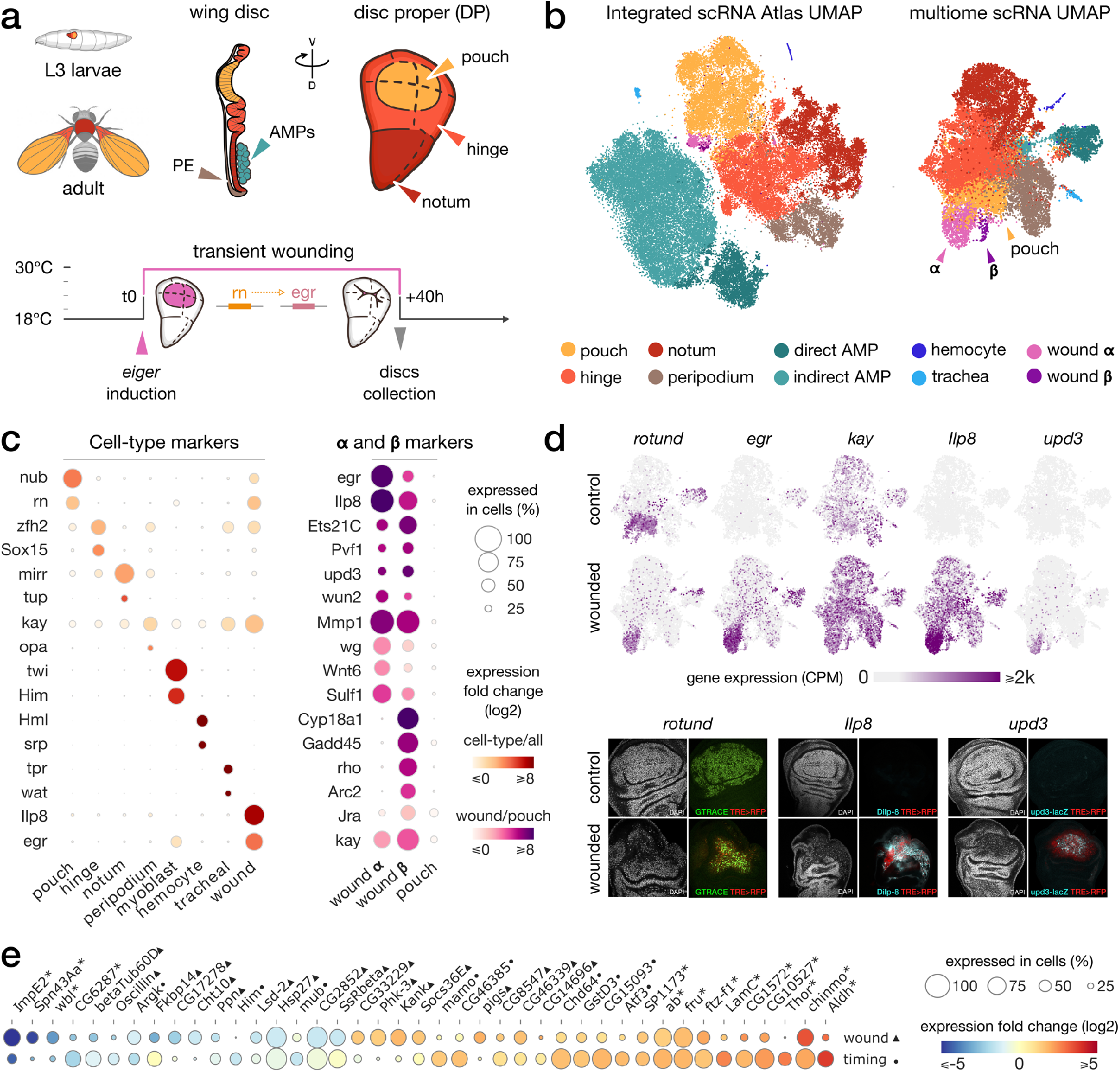
**(a) top**. Schematic of the wing imaginal disc subdomains from a Drosophila third instar larvae. The disc proper (DP) is composed of three domains and is shoehorned between the peripodial epithelium (PE) and the Adult Muscle Precursors (AMP, or myoblasts). **bottom**. Design of the wound experiment, *eiger* expression is induced via in the pouch for 40h, resulting in a localized apoptosis. **(b)** Representation of the wing disc scRNA Atlas tSNE (left) and the scRNA multiome data UMAP (right), where wound populations α and β can be detected. **(c)** Gene expression of marker genes across cell types (left) and wound subclusters in contrast to pouch (right) clusters. **(d)** Gene expression pattern of wound markers in multiome UMAP (top) and immunostaining (bottom). *Ilp8, rn* and *upd3* are found to colocalize with JNK reporter (TRE-RFP), present at the wound site. **(e)** Relative expression (log2FC) of markers of wound response (1st row, wounded/wild-type, ▴) and/or developmental timing (2nd row, 96h/120h after egg laying, •). Shared markers are marked by an asterisk (*).

## Results

### Single-cell multi-omics of the normal and wounded wing imaginal disc

To study the gene regulatory program of a wound response at single-cell resolution, we used a published genetic model that induces a sterile wound (Fortezza et al., 2016; Smith-Bolton et al., 2009)(see Methods). In this model, the expression of the TNF ligand *eiger* (*egr*) is induced in the wing pouch, where *rotund* (*rn*) is expressed, at specific times through a temperature shift regime. The expression of *eiger* causes a wound by inducing extensive local cell death (Figure 1.a). Wing discs were dissected 40 hours after the induction of *rn*-GAL4, the time point when most *rotund*-expressing cells have undergone apoptosis, and markers of wound response are observed (Herrera et al., 2013; Pérez-Garijo et al., 2013). After disc dissociation, we performed multiome experiments using 10x Genomics (combined scRNA & scATAC from the same cell), as well as on wild type control discs. This yielded 17,402 high-quality cells, with a median of 1124 detected genes per cell, and a median of 3060 unique ATAC fragments per cell (Supplementary Figure 1).

To increase the power to robustly detect cell types, and to simultaneously validate our single-cell data, we integrated our scRNA dataset with several previously published scRNA-seq datasets of wild-type wing imaginal discs, generating an integrated atlas. This wing atlas is available through SCope, along with clustering and marker gene information (Figure 1.b, Supplementary Figure 2.a, https://scope.aertslab.org/#/WingAtlas/*/welcome). The integrated atlas contains 70,230 cells from 10x Genomics, across 10 replicates (four from this work, four from (Everetts et al., 2021), one from (Bageritz et al., 2019), and one from (Deng et al., 2019)).

To annotate the cell types of the wild type disc, we compared markers from literature with significantly up-regulated gene sets obtained from each cluster (see Methods), resulting in the identification of the previously known wing disc cell types and patterning domains (Figure 1.c, Supplementary Table 1). We also confirmed our annotation by comparison with the annotations from the integrated public datasets (Supplementary Figure 2.b). The annotated clusters form a continuum of epithelial cells, from the pouch over the hinge to the notum (globally marked by the epithelial marker *grainyhead*); and separate clusters of myoblasts (marked by myogenic genes such as *twist, holes in muscle* and *zn finger homeodomain 1*), hemocytes, and tracheal cells (Figure 1b, Supplementary Figure 2.c).

One cluster was noticeably enriched for cells from the wounded disc samples, with markers linked to stress response pathways (*e*.*g*., *kayak, insulin-like peptide 8, unpaired3*, p-adj<10e^-3^, log2FC>1.7; Figure 1.c, Supplementary Figure 2.d) and localized at the wound site in the pouch domain (Figure 1.d), suggesting that this cluster represents a wound-response cell state. This cluster can furthermore be subdivided at higher resolution into two distinct cell populations (population **α** of 1,211 cells and population **β** of 94 cells, Figure 1.b-c).

Next, we analyzed the scATAC-seq part of our multiome dataset separately using cisTopic (Bravo González-Blas et al., 2019) (Supplementary Figure 2.e). Since multiome data delivers same-cell measures for RNA and ATAC, we could label the scATAC-seq based on the previously derived scRNA-seq annotations. The detected ATAC clusters, based solely on chromatin accessibility, also identify the hinge, pouch, notum, myoblast, peripodial epithelium and wound clusters (Supplementary Figure 2.f). Thus, both chromatin accessibility and gene expression independently identify normal cell types and a separate wound cell state.

### The wounded disc is delayed in its developmental timing

The proper regeneration of a damaged wing disc is tied to the introduction of a developmental delay via a reduced ecdysone signaling. This delay provides the necessary time for tissue repair before pupation (Jaszczak and Halme, 2016; Katsuyama et al., 2015; Sanchez et al., 2019). To assess whether we can detect this regulatory response in our integrated dataset, we combined cells from multiple conditions (wounded / wild-type) but also normal discs dissected at different developmental time points (96 and 120 hours after egg laying, from (Everetts et al., 2021)). By comparing the up and downregulated genes with respect to the developmental time, we found that marker genes of late time points (*e*.*g. ecdysone-inducible gene E2*) are globally down-regulated in the wounded disc samples dissected at the same time point. We furthermore found a significant overlap of down-regulated markers for both wound response and developmental timing (21%, p-adj<10e^-3^, Fisher’s exact test, Figure 1.e). We confirmed this result when restricting the analysis outside of the wound site, in the notum domain (34% shared down-regulated genes, p-adj<10e^-3^, Fisher’s exact test, Supplementary Figure 2.g). This resemblance of wounded disc cells with those from earlier stage wild-type larvae confirms that wounding triggers a global reaction across the whole disc which delays development to give more time for the wound to repair completely before metamorphosis. This delay is likely driven by *insulin-like peptide 8* (*ilp8*), a critical long-range regulator of ecdysone signaling, highly expressed in the wound cluster (Figure 1.c-d).

We additionally found genes significantly up-regulated in the entire wing disc following wounding, with no strong change between developmental time points (p-adj<10e^-3^, log2FC_wound_>1.7, log2FC_timing_<0.5, Figure 1.e). Among these wound markers, *suppressor of cytokine signaling at 36E* (*socs36E*) is a known target of the JAK/STAT and EGFR pathways (Berez et al., 2020; Kang et al., 2018). We also noticed an up-regulation of *pickled eggs* (*pigs*), a potential Notch regulator (Pines et al., 2010), in the hinge and peripodial cells of wounded discs.

### Multi-ome Gene Regulatory Network reconstruction

An important advantage of single-cell multiomics data is the power to detect regulatory interactions by synchronous changes in expression and/or accessibility across cells (Fiers et al., 2018). Here, we set out to infer enhancer-GRNs (eGRN) following a similar strategy as we recently applied to the fly brain (Janssens et al., 2022) (Figure 2.a, see Methods). First, we defined differentially accessible regions (DARs) for each cell type, including the wound cluster, using *cisTopic* (Bravo González-Blas et al., 2019). Next, these differentially accessible regions were tested for enrichment in TF binding motifs using *cisTarget* (Herrmann et al., 2012), which resulted in a list of TF-to-regions edges (i.e. regulatory links) with a significant motif hit (Figure 2.a.1). Next we determined co-variability between gene expression and accessibility of neighboring enhancer regions to generate a list of region-to-gene edges (Figure 2.a.2). We then completed the edges loop by using the TF-to-gene adjacency scores from *pySCENIC* (Figure 2.a.3) (Van de Sande et al., 2020). Lastly, we filtered for high-quality TF-to-region-to-gene interactions by keeping the leading edges of a gene set enrichment analysis (GSEA), taking the TF-to-gene scores as ranking values (Figure 2.a.4).

**Figure 2:**
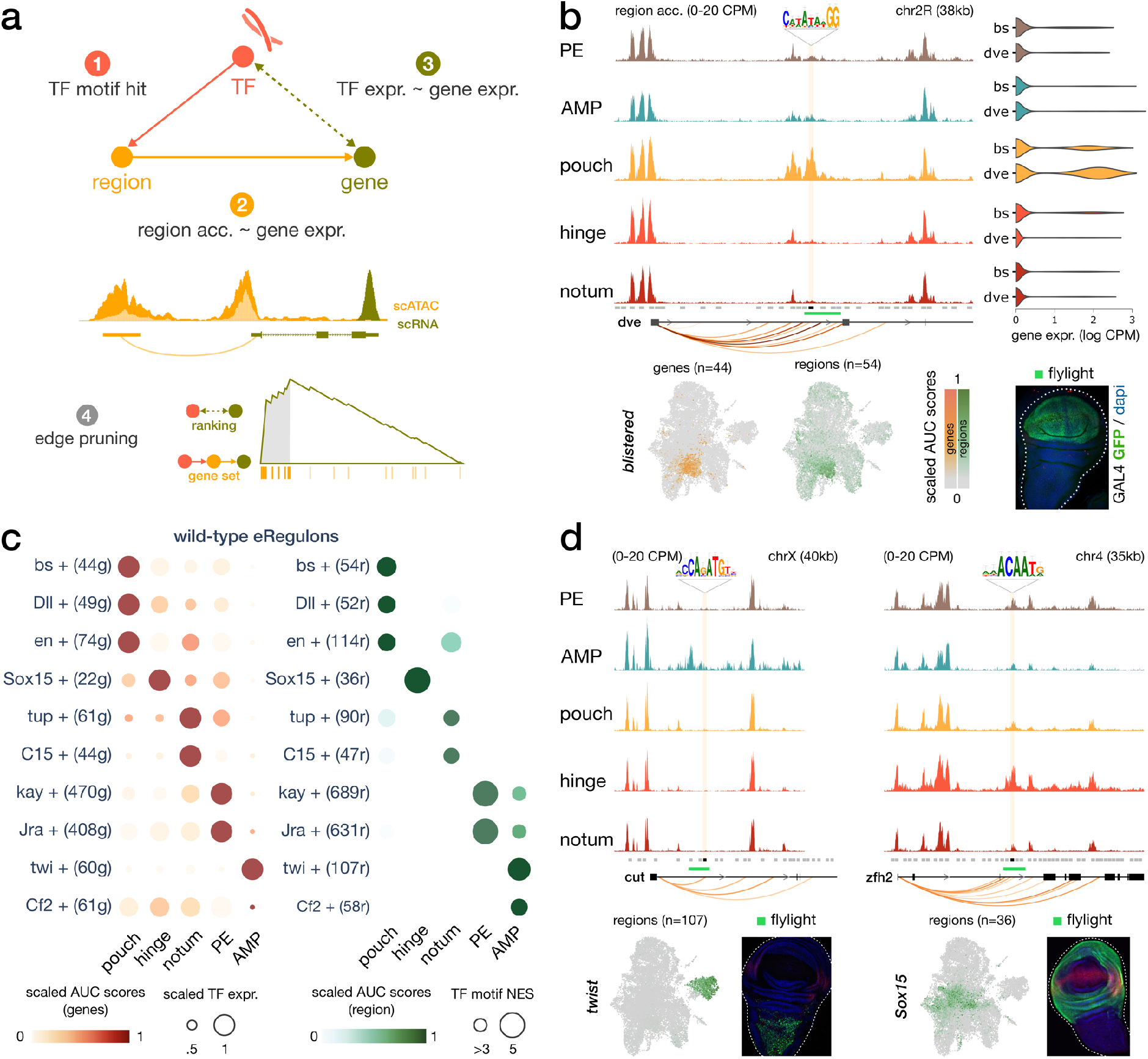
**(a)** eGRN construction is based on TF motif enrichment (1), co-variability of gene expression and chromatin accessibility (2-3), and followed by functional edge selection (4). **(b)** Aggregated wild-type chromatin accessibility signals around a target region (black rectangle) of the *blistered* eGRN. The target region comprises a *blistered* motif hit, it is significantly correlated with expression of *defective proventriculus* (*dve*, orange arcs), and it overlaps a flylight reporter (green rectangle) expressed in the pouch domain. The activities of the *blistered* TF and its associated eGRN signatures are all similarly localized in the pouch (UMAP and violin plots). **(c)** Dotplot of the average gene-based (left) and region-based (right) activity of selected eGRNs with highest cell type specificity in wild-type conditions. **(d)** Aggregated chromatin accessibility signals similar to panel **b** for the target regions of two other eGRNs, active in the myoblast (*twist*, left) and the hinge domains (*Sox15*, right).

This eGRN inference approach has the advantage of including distal enhancers, found up to 50kb away from a putative target gene. This procedure resulted in 147 high-quality eGRNs (85 activating, 62 repressing), spanning 98 TFs, with on average 54 target genes and 58 target regions (Supplementary Table 2). An example of an inferred eGRN is the TF *blistered* (*bs*), which is predicted to target the repressor *defective proventriculus* (*dve*) via multiple intronic enhancers (Figure 2.b). Using the target genes and target regions sets as proxies, we scored the activity of a TF and its subsequent domain specificity via its entire eGRN by *AUCell* (Figure 2.c, see Method). In the case of *bs*, we found the eGRN to be specifically active in the pouch, both from the gene expression and the region accessibility perspectives (Figure 2.b-c).

We can further divide our final list of eGRNs into repressor and activator categories, based on linear correlation between TF expression and accessibility. For example, we identified *mirror* (*mirr*) as a repressor TF in control wing discs, because its expression negatively correlated with accessibility of both its target genes and regions (Supplementary Figure 3.a). The repressive action of Iroquois TFs like *mirr* has already been demonstrated in Drosophila (Andreu et al., 2012; Bilioni et al., 2005).

We used the TF activity scores derived from gene expression and chromatin accessibility to compute the regulon specificity score (RSS) of each eGRN across the disc domains (Suo et al., 2018). A high RSS score indicates an enrichment for the TF signature (gene or region targets) among the top markers of a given domain. One of the strongest regulatory programs was observed in the hinge with the Sox box protein 15 (Sox15) eGRN (22 genes, 36 regions, Figure 2.c-d) where the top predicted target genes included *zn finger homeodomain 2, frizzled, dachsous* and *homothorax*. Several other well-known wing development TFs were identified, including *tailup* and *odd-paired* in the notum, *ultrabithorax* and *C15* in the peripodial epithelium and *twist* in the myoblasts (Figure 2.c-d, Supplementary Figure 3.a-c). Another interesting example is *nubbin* (*nub*), a POU/homeodomain transcription factor targeted by *vestigial* (Rodríguez et al., 2002), which is found specifically expressed in the pouch domain (Supplementary Figure 3.a). An extended list of TFs and their target genes can be queried via SCope (https://scope.aertslab.org/#/WingAtlas/*/welcome), and is provided as a Supplementary Table 2.

### The wound site shows a strong JNK and JAK/STAT eGRN activity

Both **α** and **β** wound clusters are unique to the wounded disc and share enrichment for GO terms related to wound response and paracrine signaling (p-adj<10e^-3^). They both express high levels of stress-response genes, including *ilp8* (Katsuyama et al., 2015), *matrix metalloproteinase 1* (*Mmp1*, (Harris et al., 2020)), *moladietz* (*mol*, (Khan et al., 2017)), *PDGF-and VEGF-related factor 1* (*pvf1*, (Wu et al., 2009)), and *jun-related antigen/kayak* (*jra/kay*, homologs of JUN and FOS, forming the AP-1 complex, involved in the JNK cascade (Cosolo et al., 2019)) (Figure 1.c-d, p-adj<10e^-3^, log2FC>1). The activity of the JNK pathway is further confirmed by a high and specific activity of the Jra/Kay eGRNs in the wound populations, at both gene expression and chromatin accessibility levels (Figure 3.a). We also confirm the strong involvement of JAK/STAT, as the eGRN of Signal-transducer and activator of transcription protein at 92E (Stat92E) is specifically active in the wound response clusters (Figure 3.a), in agreement with the enrichment for the Stat92E binding motif in the wound-specific accessible regions (NES=3.6) and the co-expression of the *unpaired* ligands (*upd1/2/3*, Figure 1.c).

**Figure 3:**
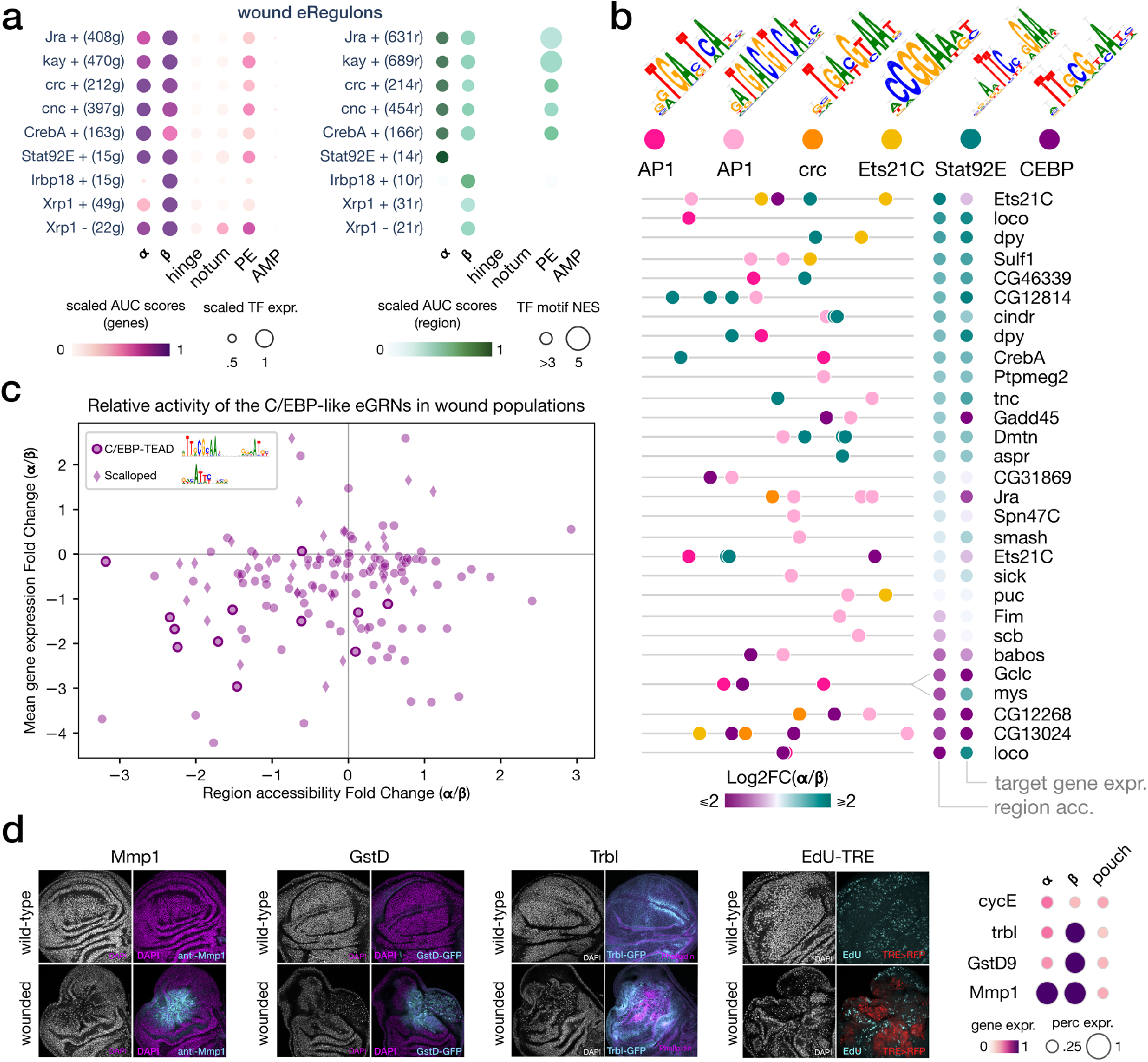
**(a)** Dotplot of the average gene-based (left) and region-based (right) activity of selected eGRNs with highest α and/or β specificity in wounded conditions. **(b)** Feature maps of six types of TF motifs on regions specifically accessible in the wound populations α and/or β. AP1 bindings are homogeneously present, while Stat92E and C/EBP motifs are specifically enriched in wound population α and β, respectively. One region targeted by both AP1 and C/EBP GRNs regulates two genes with antagonistic α/β expression patterns (mys and Gclc). **(c)** Scatterplot of the relative changes in target gene expression and chromatin accessibility for all enhancers targeted by the C/EBP eGRNS (vri, Irbp18, Xrp1, slbo). We note the presence of the CEBP-TEAD dimer motif in regions strongly upregulated in β. **(d)** Wing disc immunostaining (left) and normalized average expression of three wound marker genes and a proliferation marker (CycE, right). Both α and β marker genes are expressed and localized at the wound site.

Other transcription factors with induced eGRN activity in both wound clusters include the Cyclic-AMP response element binding protein A (CrebA, CREB3L2 ortholog, involved in resistance to infection), Cap’n’collar (Cnc, Nrf1/2 ortholog, involved in oxidative stress response), and Cryptocephal (Crc, ATF4 ortholog, involved in unfolded protein response)(Figure 3.a), three basic-leucine zipper (bZip) TFs known to be involved in stress response (Brock et al., 2017; Brown et al., 2021; Ragheb et al., 2017; Sorge et al., 2020; Sykiotis and Bohmann, 2008; Troha et al., 2018). Interestingly, we note an additional up-regulation of both Stat92E and AP-1 GRN activity in the peripodial epithelium (PE) of the wounded disc (Figure 3.a). Although the PE was not directly targeted by the genetic ablation, this domain is in close proximity with the pouch territory in wild-type wing discs (Figure 1.a). We therefore hypothesized that stress signaling is capable of local diffusion across epithelial layers. We further identified a JAK/STAT repressor *apontic* (*apt*), to be specifically expressed in the peripodial epithelium in wounded conditions (p-adj<10e^-3^, log2FC>1). This finding supports a protective role of Apontic to block a response to Stat92E proliferative signaling in the vicinity of wounded tissues (Harris et al., 2020).

Taken together, our results highlight JNK and JAK/STAT as the most prominent markers of the global wound response program, shared by the two cell populations **α** and **β**. AP1 is the largest inferred wound eGRN (470 target genes, 689 target regions) and its binding motifs (CRE and TRE variants (Fonseca et al., 2019)) are strongly enriched in differentially accessible regions from both wound populations **α** an **β** (Figure 3.b).

### Proliferative and senescence signals separate the α and β populations

In the wound samples, we expect the *rn*-expressing pouch cells to be almost entirely lost upon apoptosis from the *rn*-GAL4 induction. However, we find persistent *rn*-positive cells in the wound cluster **α** (p-adj<10e^-3^, log2FC>2), that are excluded from cluster **β** (p-adj_α/β_<10e^-3^, log2FC_α/β_>2). These cells are distinct from the normal *rn*-positive pouch cells though, as they also show stress response markers. We further observe an up-regulation of markers of tissue patterning and proliferation in cluster **α** relative to cluster **β**, like *wingless* (*wg*), *Wnt oncogene analog 6* (*Wnt6*) (Figure 1.c, p-adj_α/β_<10e^-3^, log2FC_α/β_>1) and *CyclinE* (*CycE*) (Figure 3.d, log2FC_α/β_>0.8). In this line, we also find evidence for an up-regulation of the pro-regenerative marker *Ets at 21C* (*Ets21C*) (Worley et al., 2021), with a combined enrichment of its gene expression (p-adj<10e^-3^, log2FC>8) and its binding motif in cluster **α** (*cistarget*, NES=3.35). Consistent with the higher enhancer activity of the Stat92E eGRN in wound **α** compared to **β** (Figure 3.a-b), we find a significant overlap between the **α** marker genes and a study that identified JAK/STAT as coordinating cell proliferation during wing disc regeneration (GSEA, NES=1.292)(Katsuyama et al., 2015). These results demonstrate that pro-proliferative and pro-regenerative characteristics are specific to the **α** population in the wound.

In contrast, cells from cluster **β** do not show clear proliferative nor regeneration markers. Genes found up-regulated in **β** compared to **α** include genes associated with innate immunity (*Dorsal-related immunity factor*), glutathione metabolism (*Glutamate-cysteine ligase catalytic subunit, Glutathione S transferase D9*)(p-adj_β/α_<10e^-3^, log2FC_β/α_>2), cell migration signaling (*e*.*g rho1, slow border* and *pvf1*, p-adj<10e^-3^), response to irradiation (e.g. *inverted repeat-binding protein*, p-adj<10e^-3^) and regulation of cell cycle (e.g. *growth arrest and DNA damage-inducible 45, tribbles*). Among these markers, *tribbles* (*trbl*) and *slow border cells* (*slbo*) respectively encode a known repressor and target of the JAK/STAT pathway in border cell migration (Berez et al., 2020; Dobens et al., 2021; Harris et al., 2020; Rørth et al., 2000; Starz-Gaiano et al., 2008). The cells expressing these markers are located at the center of the wounded pouch, despite their lack of pouch-specification markers (Figure 3.d). These results suggest that cluster **β** contains cells derived from the wild-type, *rn*-expressing domain, that have de-differentiated and hence no longer express wing disc markers.

In addition to their loss of wing fate, the cells from cluster **β** show interesting similarities with cellular senescence. Indeed, a recent study in wounded wing imaginal discs associate the emergence of senescent-like cells with the presence of markers of stress-response (*jra, kay*), DNA-damage response (*gadd45*), paracrine signaling (*pvf1*) and cell cycle stalling (*trbl*), as observed in our population **β** (Cosolo et al., 2019). Precisely, one marker of our cluster **β** population, *tribbles* (*trbl*), is found to be partly responsible for cell cycle stalling in wounded cells. The *slbo* expression in population **β** further corroborates the proximity with senescence-associated secretory phenotype (SASP), as *slbo* is homologous to the human C/EBPB homodimer, acting in SASP (Lee et al., 2010b, 2010a) (Reactome ID R-DME-2559582). Taken in combination with the lack of pouch markers, our results show that the population **β** corresponds to cell that lost their initial pouch fate, arrested cycling and activated a cellular senescence program.

### The senescent population is characterized by C/EBP eGRN activity

Looking at eGRN activities, we find several TFs with a strong specificity score for the senescent population (β), namely the heterodimer partners *inverted repeat binding protein 18 kDa* (*irbp18*) and *xrp1, slbo, vrille* (*vri*) and *sox box protein 14* (*sox14*) (top 6% RSS, Figure 3.a). Among them, the two bZip TFs *irbp18* and *vri* are significantly upregulated in the senescent population compared to the proliferative population (**α**) (p-adj<2*10e^-3^, log2FC_β***/***α_>2.4) and share similarities in their TF binding motifs with the senescence-associated C/EBPB dimers (Figure 3.b)(Blanco et al., 2020). The Irbp18 eGRN (extended category, see method) is composed of 149 predicted target regions (80 wound-specific and 11 senescent-specific regions) and 116 target genes (52 markers of senescent population, p-adj<5*10e^-3^, log2FC>1.5) including *slbo, vri, ets at 98B* (*ets98B*), *socs36E, head involution defective* (*hid*), *growth arrest and DNA damage-inducible 45* (*gadd45*) and *p53*. The presence of these TFs in the senescent cluster (**β**) is further validated by the significant enrichment for C/EBPB-like binding motifs in senescent-specific peaks (NES = 5.25, Figure 3.b-c) generated using MACS2 *bdgdiff* (Y. Zhang et al., 2008).

Our eGRN approach has an important limitation, namely that TFs can be identified only if their expression co-varies with chromatin accessibility and target gene expression. A key TF that remains undetected in the eGRN approach is Scalloped (Sd). This mammalian TEAD homolog is the effector TF of the Hippo signaling, a pro-proliferative pathway known to be activated during tissue growth and regeneration (L. Zhang et al., 2008). Still, we find a significant enrichment for the Sd motif in the senescent-specific accessible regions (NES = 4.89). Moreover, the top enriched motif in these senescent-specific regions is a C/EBP-TEAD dimer motif (NES = 7.48, Figure 3.c), notably found in the enhancer of CG13024, a strong marker of both senescent and *ras*^V12^ *scrib*^-/-^ cells (Supplementary Figure 4, see next section).

Our results suggest that the eGRN activity of C/EBP orthologs, such as the Irbp18-Xrp1 heterodimer, is a strong marker of cellular senescence. Given that these eGRNs target multiple markers of cell migration, cell cycle stalling and DNA damage response, we hypothesize that C/EBP signaling may be responsible for establishing a state of cellular senescence in the wound vicinity. By transiently escaping cell death, such cells play an essential role in stress response induction, and may serve as a “flagship” population to lead the regenerative cells toward wound closure (Cosolo et al., 2019; Kozyrska et al., 2022). The list of all significant marker genes for both wound populations is available in the Supplementary Table 3.

### Shared regulatory programs between wound response and cancer

Multiple pathways associated with senescence, such as JNK and JAK/STAT, have been found to be associated with tumorigenesis when their activity becomes unrestricted (Atkins et al., 2016; Cosolo et al., 2019; Davie et al., 2015; Fortezza et al., 2016; Külshammer et al., 2015; Pinal et al., 2019; Uhlirova and Bohmann, 2006). One oncogenic system widely used is the *ras*^V12^*scrib*^-/-^ tumor model (Brumby and Richardson, 2003), where the *scribble*^-/-^ (*scrib*^-/-^) mutation causes epithelial cells to lose polarity, while *raspberry*^V12^ (*ras*^V12^) prevents them from being outcompeted (Figure 4.a). The constant stress caused by tissue disorganization, combined with the lack of apoptosis, leads to an overgrown population of cells in the eye-antennal disc (Atkins et al., 2016; Davie et al., 2015), described as *aberrant senescence* (Cosolo et al., 2019; Rhinn et al., 2019). In order to compare the senescent state (**β**) observed in our transient wound model with tumor overgrowth, we performed scRNA-seq of *ras*^V12^*scrib*^*/-*^ eye-antennal discs (6,717 cells).

**Figure 4:**
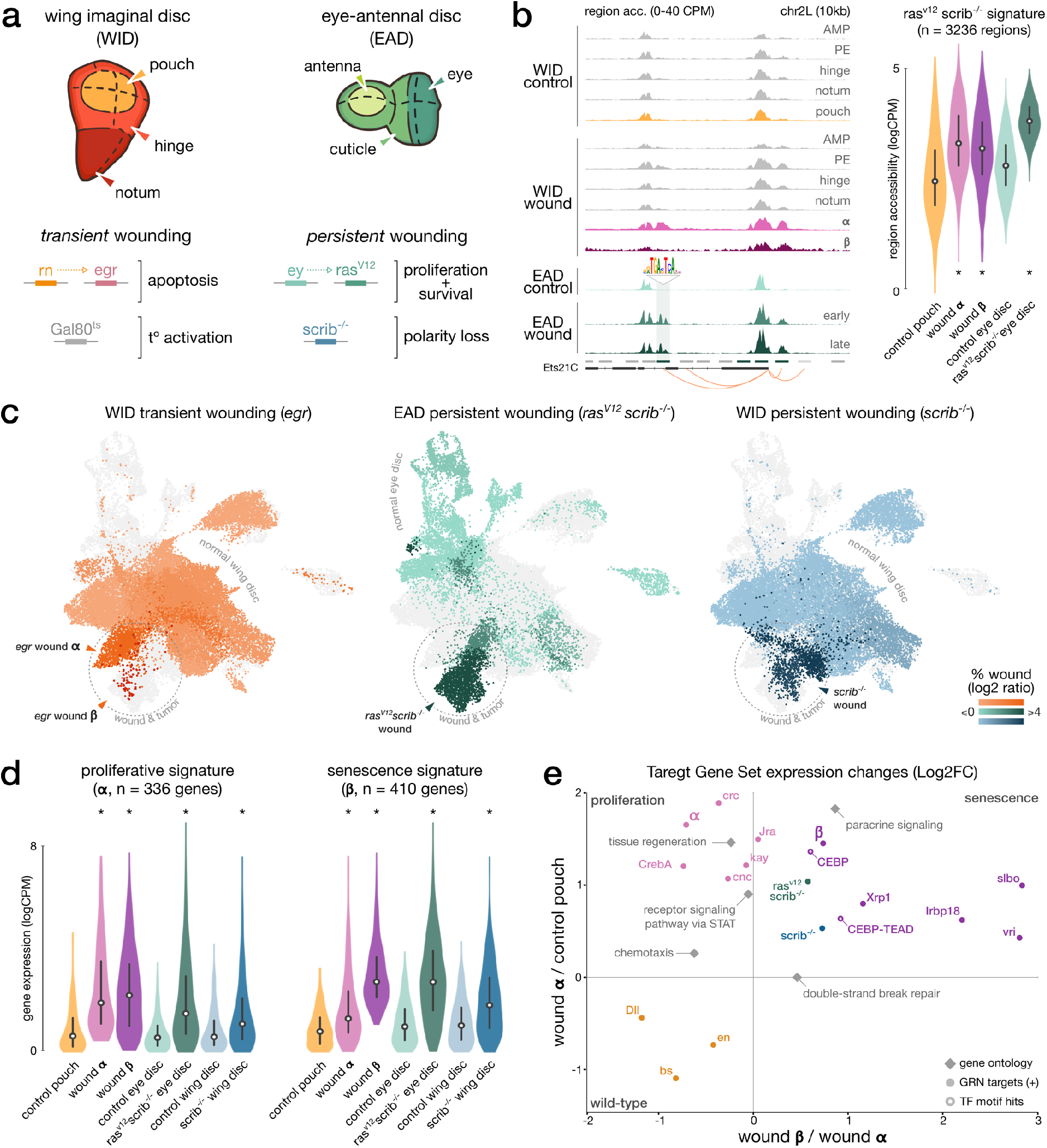
**(a) top**. Schematic of the imaginal wing and eye-antennal discs. **bottom**. Genetic constructs used to trigger localized, transient or persistent wounding. **(b) left**. Aggregated (WID) and bulk (EAD) chromatin accessibility signals in wild-type and both wound conditions at Ets21C loci, including an AP-1 target region. **right**. Distribution of ATAC measures in *ras*^*V12*^*scrib*^*-/-*^ region signature, asterisks denotes significant enrichment compared to wild-type. **(c)** Integrated wound atlas UMAP. Dark color clusters are enriched in wound cells. We detect our populations **α** and **β** from the *egr* wound and two additional *ras*^*V12*^*scrib*^*-/-*^ and *scrib*^*-/-*^ wound populations. **(d)** Distribution of RNA measures in wound **α** and **β** gene signature, asterisks denotes significant enrichment compared to wild-type. **(e)** Scatter plot of the mean expression fold-change of gene sets extracted from eGRN, GO, marker genes (*ras*^*V12*^*scrib*^*-/-*^ and *scrib*^*-/-*^) and β-specific regions with CEBP or CEBP-TEAD motifs hits (target genes inferred from the CEBP eGRNS). Fold-changes are contrasting gene expression changes between x) wound population αand **β** and y) between wild-type pouch domain and wound α. All elements are significantly up-or down-regulated in at least one axis (p-adj<5*10e^-3^, paired t-test, Bonferroni corrected).

We combined this dataset with publicly available scRNA-seq data on wild-type eye-antennal discs (Bravo González-Blas et al., 2020) (4,830 cells), scRNA-seq data on 14-day *scrib*^-/-^ imaginal wing discs (Deng et al., 2019; Ji et al., 2019)(7,554 cells) and bulk ATAC-seq data from *ras*^V12^*scrib*^-/-^ tumors (Davie et al., 2015). Altogether, these datasets allowed us to compare wound response programs across different model systems (eye and wing discs), regulatory layers (RNA and ATAC), and conditions (following wound induction and during persistent oncogenic induction).

We first characterized regions specifically accessible in the *ras*^V12^*scrib*^-/-^ model compared to wild type eye discs (Figure 4.b). In agreement with Davie *et al*., this signature is enriched for the binding motifs of Stat92E (NES = 4.19) and AP-1 (NES = 7.17). Looking at pseudo-bulk ATAC aggregate wing disc data, we noted a significant enrichment for the *ras*^V12^*scrib*^-/-^ signature regions for both proliferative (α) and senescent (**β**) wound populations in contrast to wild-type pouch cells (p-adj<10e^-3^, one-tailed t-test, Figure 4.b). In line with the observed AP-1 and Stat92E activity in wounded peripodial epithelium, we also find the *ras*^V12^*scrib*^-/-^ signature enriched in this domain (Supplementary Figure 5.a). These similar chromatin accessibility profiles between wound and tumor populations suggest at least part of the *ras*^V12^*scrib*^-/-^ cells also activate the senescence and proliferative programs.

We further investigated the similarities with oncogenic induction at the gene expression level by integrating all single-cell RNA datasets into a combined, batch-corrected, analysis (44,759 cells, see Methods). For each oncogenic model (*ras*^V12^*scrib*^-/-^ and *scrib*^-/-^), we detect a tumor-specific cluster, mostly composed of cells originating from samples with induced tumorigenesis (Figure 4.c). This analysis reveals a co-clustering of the senescent wound cells (β) with *ras*^V12^*scrib*^-/-^ and *scrib*^-/-^ cells (Figure 4.c). We confirm the similarity in gene expression patterns between oncogenic models and both wound populations by computing the activity of the proliferative and senescent signatures (**α** and **β** relative to control pouch) in wild-type and oncogenic conditions (Figure 4.d). We find both signatures to be upregulated, with the strongest increase in gene expression between wild-type and the two oncogenic models for the senescent signature (p-adj<10e^-3^, one-tailed t-test, Figure 4.d). We find a similar result for signatures contrasting the proliferative and senescent populations against each other (α vs. β), further supporting the closest resemblance between the chronic oncogenic response and senescent population (Figure 4.e, Supplementary Figure 5.b). This result suggests that the tumor and senescent populations all de-differentiated from their original state (eye or wing discs) toward a shared wound-response program. Yet, unlike the wound response, we do not observe two distinct proliferative-like and senescent-like cell populations in tumors. On the contrary, the tumor population is homogeneous, which suggests that tumor cells may simultaneously activate both pro-proliferative and pro-senescence programs in an aberrant combination, undetected in a genetically wild-type context.

## Discussion

In this study, we used several wound paradigms from Drosophila imaginal discs, both transient (following normal wounding) and persistent (continuous wound induction during tumor formation), to describe the mechanisms coordinating stress response. We constructed eGRNs from single-cell ATAC and RNA data, assigning TFs to their inferred regulatory target genes and target enhancer regions. Using this new eGRN viewpoint, we found two distinct populations respectively displaying proliferative (**α**) and senescent (**β**) properties. Both populations share a common wound response program, comprising JNK and JAK/STAT signaling. Yet, the senescent cells display an additional activation of DNA damage, paracrine signaling and stress mitigation pathways (glutathione metabolism, FC_α/β_>1.8) that is distinct from the proliferative cells. In agreement with a possible de-differentiation process, TFs associated with pouch fate specification are also down-regulated in senescent cells (*Distal-less, blistered, engrailed*). Concurrently, the senescence markers co-vary with C/EBP-like eGRN activity and the oncogenic signatures (*ras*^V12^*scrib*^-/-^ and *scrib*^-/-^, top 100 target genes, Figure 4.e). Taken together, these results highlight a common response program, shared by all wound populations and *ras*^V12^*scrib*^-/-^ cancer cells (JNK, JAK/STAT), and an additional senescent-like program driven by C/EBP-like TFs that is shared between the senescent population and the tumorigenesis paradigms. To our knowledge, this study presents the first depiction of the senescence phenotype at the single cell multiomic level (scRNA + scATAC). We believe our characterized gene and enhancer signatures will help pave the way toward a universal marker set of senescence, which is currently outstanding (Cohn et al., 2022; Gorgoulis et al., 2019). Using these single-cell senescent profiles along with their eGRN signatures, we can already show here a close proximity between the senescence phenotype and oncogenic programs, mostly due to the shared upregulation of C/EBP transcription factors activity.

The persistent wound paradigm (e.g. *ras*^V12^*scrib*^-/-^) causes uncontrolled neoplastic growth and has previously been referred to as a tumor model (Dillard et al., 2021; Mirzoyan et al., 2019). In this study, we found tumor programs to be highly similar to the senescent-like cell population emerging following a transient wound. These findings corroborate the growing set of evidence suggesting a large overlap between the hallmarks of tumor and wound response (MacCarthy-Morrogh and Martin, 2020). From this perspective, we can see a tumor paradigm as a wound that never heals, where the failure to arrest cell proliferation would prevent the damaged cells to fully resorb.

The senescent phenotype has been extensively studied, notably in the context of wound repair (Demaria et al., 2014; Rhinn et al., 2019; Wilkinson and Hardman, 2020). It is defined as a state of arrested cell proliferation, combined with a strong paracrine signaling activity (SASP) promoting stress-response signaling and partly mediated by CEBP/B activation. Recent studies further suggest a guiding role, where senescent cells lead the collective migration of proliferative cells toward wound closure, where they are then discarded through mechanical competition (Kozyrska et al., 2022).

The mechanism mediating the clearance of senescent cells is critical to prevent the persistence of stress signaling, as observed in tumor models (Baumgartner et al., 2021; Kozyrska et al., 2022; Pinal et al., 2019). In line with the hypothesis of competition-mediated clearance of senescent cells (Kozyrska et al., 2022), recent studies developed the importance of C/EBP in stress response and cell competition, although no clear consensus currently exists regarding its type of action (promoting or inhibiting growth and survival) (Baillon et al., 2018; Blanco et al., 2020; Boulan et al., 2019; Brown et al., 2021; Huggins et al., 2015; Lee et al., 2018; Logeay et al., 2022). Given the ambivalent needs for senescent cells to firstly survive to a strong apoptotic signal (ablation) and to secondly get discarded through competition (closure), we believe that further investigations the C/EBP-mediated regulation of apoptosis susceptibility will be of high interest. Following this hypothesis, failure to arrest SASP signaling through senescent cell clearance would promote overproliferation in persistent oncogenic induction. This idea of competition-mediated clearance is supported by several studies showing that polarity sensing mutants (e.g. *scrib*^*-/-*^) would overproliferate when present as a homogeneous population, but would be cleared within a crowd of wild-type cells (Ballesteros-Arias et al., 2014; Brumby and Richardson, 2003; Pinal et al., 2019).

The mechanism governing the sporadic emergence and persistence of senescent cells at the wound site is still unclear. Previously it was shown that the DNA damage responsive TF p53 and cell cycle stalling cooperate to trigger cellular senescence (Kozyrska et al., 2022). Such stalling has also been reported in the Drosophila wound model, where it is mediated by AP-1 and involves *tribbles* (Cosolo et al., 2019), two markers of our senescent population. The authors further highlight that JNK activity is protecting the damaged, senescent cells from competition as long as they do not resume cycling. Two other studies have also demonstrated the role of a positive feedback loop via *moladietz* (marker of our proliferative population **α**) to sustain JNK activity in the senescent cells after wounding (Khan et al., 2017; Pinal et al., 2018). Additional studies suggest the action of microRNAs (Bilak et al., 2014) and transposable elements (TE) (Azpiazu and Morata, 2022; Colombo et al., 2018) to induce cell senescence via chromatin remodeling pathways. In this line, we find specific expression of the retrotransposons TAHRE and Invader4 in the wound populations (p-adj<10e^-3^, log2FC>1, Supplementary Figure 5.c), although, further investigation would be necessary to fully characterize this TE-mediated response.

We do not detect a strong signal for the signature of several TFs that have been previously suggested to play a role in the wound response, such as p53. Indeed, we do not find evidence for the activity of p53 eGRN at the chromatin level (Jacobs et al., 2018), even in the senescent population where it is present as a C/EBP target gene (Supplementary Figure 4.a). This is also not the case for *Grainyhead*, which is involved in ERK signaling and epithelial barrier repair (Mace et al., 2005), but is not up-regulated in the wound response state (Supplementary Figure 5.d). The Hippo pathway (pro-proliferative and pro-invasive in *ras*^V12^*scrib*^-/-^) is also not detected as a wound signature in our *rn*-*egr* model, although we find an enrichment of its effector’s motif, Scalloped, in the senescent wound population. The presence of *sd* is consistent with the reported cooperation of its human ortholog, TEAD, with AP-1 in invasive mesenchymal-like cells (Verfaillie et al., 2015; Zanconato et al., 2015).

Based on the eGRN signatures, our results indicate that our wound population **β**, marked by high paracrine and DNA-damage signaling, corresponds to a senescent cell type. Our data supports a model in which these senescent cells are capable of surviving apoptotic signals by cell-cycle stalling, and then further guide the migration and growth of the pro-regenerative population (**α**) via its strong secretory phenotype, notably the JNK pathway (i.e. presence of Wg and Upd family ligands (La Marca and Richardson, 2020)). The close similarity of senescent cells with *ras*^V12^*scrib*^-/-^ cells suggest that they share the same regulatory wound programs, highlighting the fact that mechanisms present in oncogenic induction may already be active in a genetically wild-type context. Yet, instead of two distinct populations of **α**-like and **β**-like cells in *ras*^V12^ *scrib*^-/-^ model, we find an activation of both signatures in the same cells, which might suggest an aberrant combination of both senescence and proliferation programs in tumorigenesis.

Given the preponderance of the C/EBP eGRNs we observe in tumor paradigms, we believe that further characterisation of their signaling, especially through TF cooperation, will help to study the biology of cancer. In this line, multiple bZip TFs present in our study are conserved in the mammalian inflammatory pathways and are capable of forming heterodimers (Blanco et al., 2020; Deppmann et al., 2006, 2006; Huggins et al., 2015; Newton and Dixit, 2012; Shokri et al., 2019). Notably, a recent study in YAP/TAZ-bound regions upregulated in breast cancer has detected an increased binding co-occurrence for the orthologs of our key regulatory TFs : AP-1, STAT3, C/EBP and TEAD (Jra, kay, Stat92E, Sd, Irbp18, slbo)(L. He et al., 2021).

## Materials and Methods

### Fly genotypes

The following fly stocks were used for ras^v12^, scrib^-/-^ experiments : y,w,eyFlp; act>y+>Gal4, UAS-GFP/UAS-RasV12; FRT82 tub-Gal80/FRT82 scrib-,e. The fly stocks for *egr* experiments are w^1118^; +; rn-Gal4, UASeiger, tub-Gal80^ts^/TM6B.

### Genetic cell ablation using Gal4/UAS/Gal80ts

To induce expression of *egr*, experiments were carried out as described in (Cosolo et al., 2019; Fortezza et al., 2016; Smith-Bolton et al., 2009) with few modifications. Briefly, larvae of genotype *rn-GAL4, tub-GAL80*^*ts*^ and carrying the desired *UAS-*transgenes were staged with a 6h egg collection and raised at 18°C at a density of 50 larvae/vial. Overexpression of the TNF ligand *egr* transgene was induced by shifting the temperature to 30°C for 24h at day 7 after egg deposition and larvae were dissected right after. Imaginal wing discs were collected from wandering third instar larvae in PBS and flash-frozen in liquid nitrogen.

### Neoplastic growth induction using ras^v12^, scrib^-/-^

Flies were raised at 25°C on a yeast-based medium under a 12h-12h day-night light cycle. Ras^v12^, scrib^-/-^ early eye-antennal discs were dissected from wandering third instar larvae (day 6-7) in PBS. Ras^v12^, scrib^-/-^ late were collected four days after larvae began wandering (day 10-11); this is possible because ras^v12^, scrib^-/-^ do not pupate, but can persist more than one week in a prolonged larval stage.

### Immunohistochemistry

Wing discs from third instar larvae were dissected and fixed for 15 min at room temperature in 4% paraformaldehyde in PBS. Washing steps were performed in PBS containing 0.1% TritonX-100 (PBT). Discs were then incubated with primary antibodies in PBT, gently mixing overnight at 4°C. Tissues were counterstained with DAPI (0.25 ng/µl, Sigma, D9542), Phalloidin-Alexa Fluor 488/647 (1:100, Life Technologies) or Phalloidin-conjugated TRITC (1:400, Sigma) during incubation with cross-absorbed secondary antibodies coupled to Alexa Fluorophores (Invitrogen or Abcam) at room temperature for 2h. Tissues were mounted using SlowFade Gold Antifade (Invitrogen, S36936). Whenever possible, experimental and control discs were processed in the same vial and mounted on the same slides to ensure comparability in staining between different genotypes. Images were acquired using the Leica TCS SP8 Microscope, using the same confocal settings and processed using tools in Fiji.

### Sample and library preparation for single-cell gene expression

#### Sample preparation

Eye-antennal discs or wing discs were dissected and transferred to a tube containing 100 µl ice cold PBS. After centrifugation at 800 g for 5 min, the supernatant was replaced by 50 µl of dispase (3 mg/ml, Sigma-Aldrich_D4818-2mg) and 75 µl collagenase I (100 mg/ml, Invitrogen_17100-017). Discs were dissociated at 25°C in a Thermoshaker (Grant Bio PCMT) for 45 min at 25°C, 500 rpm. The enzymatic reaction was reinforced by pipette mixing every 15 min. Cells were washed with 1 ml ice cold PBS and resuspended in 400 µl PBS supplemented with 0.04% BSA. Cell suspensions were passed through a 10 µM pluriStrainer (ImTec Diagnostics-435001050). Cell viability and concentration were assessed by the LUNA-FL Dual Fluorescence Cell Counter.

#### Library preparation

Single-cell libraries were generated using the 10x Chromium Single-Cell Instrument and Single Cell 3’ Gene Expression (GEX) kit according to the manufacturer’s protocol. Briefly, single cells from eye-antennal discs or wing discs were suspended in 0.04% BSA-PBS. After generation of nanoliter-scale Gel bead-in-emulsions (GEMs), GEMs were reverse transcribed in a C1000 Touch Thermal Cycler (Bio Rad) programmed at 53°C for 45 min, 85°C for 5 min, and hold at 4°C. After reverse transcription, single-cell droplets were broken and the single-strand cDNA was isolated and cleaned with Cleanup Mix containing DynaBeads (Thermo Fisher Scientific). cDNA was then amplified by PCR: 98°C for 3 min; 12 cycles of 98°C for 15 s, 67°C for 20 s, 72°C for 1 min; 72°C for 1 min; and hold at 4°C. Subsequently, the amplified cDNA was fragmented, end-repaired, A-tailed and index adaptor ligated, with SPRIselect cleanup in between steps. The final gene expression library was amplified by PCR: 98°C for 45 s; 14 cycles of 98°C for 20 s, 54°C for 30 s, 72°C for 20 s; 72°C for 1 min; and hold at 4°C. The sequencing-ready library was cleaned up with SPRIselect beads.

#### Sequencing

Before sequencing, the fragment size of every library was analysed using the Bioanalyzer high-sensitivity chip. All 10x GEX libraries were sequenced HiSeq4000 or NovaSeq6000 instruments (Illumina) with the following sequencing parameters: 26 bp read 1 – 8 bp index 1 (i7) – 98 or 75 bp read 2.

### Sample and library preparation for 10x single-cell multiome ATAC and gene expression

#### Sample preparation

Control and wounded wing discs were dissected and transferred to a tube containing ice cold PBS. PBS was removed by centrifugation, tissues were flash frozen in liquid nitrogen and stored at -80°C. The following procedure was followed to extract the nuclei from the wing discs: resuspension in 500 µl nuclei lysis buffer comprising 10 mM Tris-HCl pH 7.4, 10 mM NaCl, 3 mM MgCl2, 0.1% Tween-20, 0.1% Nonidet P40, 0.01% Digitonin, 1% BSA, 1 mM dithiothreitol and 1U/µl RNasin ribonuclease inhibitor (Promega) in nuclease-free water, incubation on ice for 5 min, transfer to a dounce tissue grinder tube (Merck), 25 strokes with pestle A, incubation on ice for 10 min, 25 strokes with pestle B. The lysis was stopped by added 1 ml of wash buffer composed of 10 mM Tris-HCl pH 7.4, 10 mM NaCl, 3 mM MgCl2, 0.1% Tween 20, 1% BSA, 1 mM dithiothreitol and 1U/µl RNasin ribonuclease inhibitor in nuclease-free water. Nuclei were pelleted by centrifugation at 800 g for 5 min at 4°C and resuspended in a 1x nuclei buffer (10x Genomics) supplemented with 1 mM dithiothreitol and 1U/µl RNasin ribonuclease inhibitor. Nuclei suspensions were passed through a 40 µm Flowmi filter (VWR Bel-Art SP Scienceware). Nuclei concentration was assessed using the LUNA-FL Dual Fluorescence Cell Counter.

#### Library preparation

Single-cell libraries were generated using the 10x Chromium Single-Cell Instrument and NextGEM Single Cell Multiome ATAC + Gene Expression kit (10x Genomics) according to the manufacturer’s protocol. In brief, the single nuclei of wing discs were incubated for 60 min at 37°C with a transposase that fragments the DNA in open regions of the chromatin and adds adapter sequences to the ends of the DNA fragments. After generation of nanolitre-scale gel bead-in-emulsions (GEMs), GEMs were incubated in a C1000 Touch Thermal Cycler (Bio-Rad) under the following program: 37°C for 45 min, 25°C for 30 min and hold at 4°C. Incubation of the GEMs produced 10x barcoded DNA from the transposed DNA (for ATAC) and 10x barcoded, full-length cDNA from poly-adenylated mRNA (for GEX). This was followed by a quenching step that stopped the reaction. After quenching, single-cell droplets were broken and the transposed DNA and full-length cDNA were isolated using Cleanup Mix containing Silane Dynabeads. To fill gaps and to generate sufficient mass for library construction, the transposed DNA and cDNA were amplified via PCR: 72°C for 5 min; 98°C for 3 min; 7 cycles of 98°C for 20 s, 63°C for 30 s, 72°C for 1 min; 72°C for 1 min; and hold at 4°C. The pre-amplified product was used as input for both ATAC library construction and cDNA amplification for gene expression library construction. Illumina P7 sequence and a sample index were added to the single-strand DNA during ATAC library construction via PCR: 98°C for 45 s; 7-9 cycles of 98°C for 20 s, 67°C for 30 s, 72°C for 20 s; 72°C for 1 min; and hold at 4°C. The sequencing-ready ATAC library was cleaned up with SPRIselect beads (Beckman Coulter). Barcoded, full-length pre-amplified cDNA was further amplified via PCR: 98°C for 3 min; 6-9 cycles of 98°C for 15 s, 63°C for 20 s, 72°C for 1 min; 72°C for 1 min; and hold at 4°C. Subsequently, the amplified cDNA was fragmented, end-repaired, A-tailed and index adaptor ligated, with SPRIselect cleanup in between steps. The final gene expression library was amplified by PCR: 98°C for 45 s; 5-16 cycles of 98°C for 20 s, 54°C for 30 s, 72°C for 20 s. 72°C for 1 min; and hold at 4°C. The sequencing-ready GEX library was cleaned up with SPRIselect beads.

#### Sequencing

Before sequencing, the fragment size of every library was analysed using the Bioanalyzer high-sensitivity chip. All 10x Multiome ATAC libraries were sequenced on NovaSeq6000 instruments (Illumina) with the following sequencing parameters: 50 bp read 1 – 8 bp index 1 (i7) – 16 bp index 2 (i5) - 49 bp read 2. All 10x Multiome GEX libraries were sequenced on NovaSeq6000 instruments with the following sequencing parameters: 28 bp read 1 – 10 bp index 1 (i7) – 10 bp index 2 (i5) – 75 bp read 2.

### scATAC and scRNA read mapping

All the analyses performed in this study are using the *Drosophila melanogaster* r6.35 (dm6) reference sequence and annotations. All single-cell datasets were (re)processed using CellRangerARC/1.0.1 and CellRanger/5.0.1 with default parameters, for multiome and scRNA respectively.

### scRNA analysis

We analyzed the entire dataset with an in-house developed NextFlow pipeline, named VSN (Flerin et al., 2021). VSN performs a standard scRNA analysis with *best-practices* workflow (“Current best practices in single-cell RNA-seq analysis,” 2019), including the doublet filtering using *scrublet* (Wolock et al., 2019) and the correction for batch effects between the runs using *Harmony* (Korsunsky et al., 2019) (Supplementary Figure 2.a). scRNA data were processed with default parameters : minimum of 200 detected genes per cell, maximum of 15% signal of mitochondrial origin, minimum of 3 cells expressing any targeted genes. We additionally introduced the quality-check step from (Everetts et al., 2021) to filter for low-quality cell abundance in the integrative analysis combining our runs together with public datasets. This filtering step consists in removing cell clusters with a mean number of detected genes lower than one standard deviation below the global average. The clustering, UMAP and tSNE resulting from the integrative analysis were computed on 50 principal components (PCs, selected with the *pcacv* module (Varmuza and Filzmoser, 2009)) and a leiden resolutions of 0.5 and 1.4 for the wild-type and wound (*egr, ras*^*V12*^, *scrib*^*-/-*^) atlas integrations respectively. For the multiome-only analysis, clustering and wild-type annotations were derived from the wild-type integrated analysis. UMAP and tSNE were computed on 50 PCs. Detection of the wound sub-populations **α** and **β** was done on the same PC space, at a leiden resolution of 1.3. Marker genes were detected on log-normalized counts using the standard *scanpy rank_genes_groups* module with default or more stringent parameters (Wolf et al., 2018)(method = *t-test_overestim_var*, detailed values in the result section). Analysis of Transposable Element signal was done on the multiome data using scTE (J. He et al., 2021) with identical preprocessing steps.

### scATAC analysis

The analysis of chromatin profiles obtained from our three multiome runs was done using cisTopic (Bravo González-Blas et al., 2019). One critical step of this analysis is to confidently identify the *cis*-regulatory regions active in our system. To do so, we generated pseudo-bulk ATAC tracks by aggregating the sequencing outputs from cells with identical scRNA annotation together. For each track, we then define peaks as 500bp regions centered around MACS2-called summits (Y. Zhang et al., 2008). The consensus peak set (CPS) of 41,387 regions, additionally filtered for repeat loci, is constructed via the Iterative Overlap Peak Merging Procedure (Corces et al., 2018). In addition to the scRNA-based cell filtering (see scRNA analysis section), we filter the cells for three scATAC-based metrics : minimum number of fragment per cell of log_10_(3.5), minimum fraction of reads in CPS of 60%, minimum TSS enrichment of 2. Using the filter cell and the CPS to generate an input count matrix, we run cisTopic LDA and select the model with best topic coherence (n_topic = 76, Supplementary Figure 2.e). We use the Harmony-corrected topic dimensions to compute tSNE and UMAP embeddings (Supplementary Figure 2.f), which recapitulate the main scRNA cell types, despite a noticeable mixing of some clusters (e.g. direct and indirect AMPs are mixed and will be merged as a single annotation in the following analyses). The imputed accessibility scores are computed from the 76-topic model per cell, for each detected enhancer region. This probability matrix serves as a proxy for chromatin accessibility in the GRN inference analysis, while limiting the impact of dropout values (see method). The Differentially Accessible Regions (DARs) are computed for each cell type and conditions using a Wilcoxon rank-sum test (p-adj<5*10e^-3^, log2FC>1.5, mean 1770 DARs per cell type).

### (pseudo)bulk ATAC analysis

The wound population **β** does not form a clear cluster in the scATAC embeddings (Supplementary Figure 2.d), which might indicate a non-optimal detection of their accessibility profile in the cistopic model (potentially due to low cell number). In order to maximize the detection of the accessibility profiles of wound population **α** and **β**, we took advantage of the ATAC pseudo-bulk tracks generated previously and directly computed the differentially accessible regions between **α** and **β** using the bdgdiff module from MACS2. We find 3,990 regions accessible in both populations, 338 **α**-specific and 173 **β**-specific DARs (log-likelihood ratio>1.5, maximum gap=150, minimum length=300). The feature maps presented in Figure 2.b and Supplementary Figure 4.a were obtained by scoring a group of six TF motifs on each of these regions using cluster-buster (Frith et al., 2003) with a minimum motif score of 6. Detailed region coordinates along with motif hit information can be found in the Supplementary Table 4.

The bulk ATAC data from wild-type and *ras*^*V12*^*scrib*^*-/-*^ eye-antennal discs was reprocessed similarly to the original study (Davie et al., 2015) with updated reference genome (dm6). Reads were mapped with Bowtie2, regions were called using MACS2 and DARs were detected via DESeq2. We obtain a signature of 4,547 regions significantly more accessible in *ras*^*V12*^,*scrib*^*-/-*^ conditions compared to wild-type (p-adj<5*10e^-3^, log2FC>1.5). This signature overlaps 3,236 regions from the CPS.

### eGRN inference

The computational pipeline used to infer eGRN is principally based on previous works from (Bravo González-Blas et al., 2020) and (Janssens et al., 2022) and can be summarized in three steps schematised in Figure 2.a. In the first step, we apply *cistarget* to each of the DARs computed in the scATAC analysis (wild-type and wound, see method) as well as the pseudo-bulk contrast between wound populations **α** and **β**. We retain TF hits in each region of a set of DARs if the TF motif is overall significantly enriched in the set (NES>3, rank threshold=0.05, AUC threshold=0.005, motif similarity FDR=0.001, orthologous identity threshold=0.6) and the motif is actually detected as a hit in the region of interest. The scoring database used for this step corresponds to the cisTarget motifs collection v9 (28799 motifs), specifically re-scored on the CPS from this study (instead of the standard 131,324 candidate regulatory regions for fly i-cisTarget). To avoid issues with redundant motifs within the database (e.g. 14 motifs are associated with human JUND), we grouped similar motifs into a merged consensus prior to scoring. The final set of regions targeted by a specific TF is called a TF’s cistrome. We obtained two categories of cistromes for each TF : a direct cistrome, solely retaining results from motifs with direct TF annotation, and an extended cistrome, including enrichment results from motifs passing the similarity or orthology threshold.

In the second step, we use the log-normalized gene expression and imputed accessibility signal from our multiome runs as input (for preprocessing, see method). Using both views, we can calculate region-to-gene relationship using a non-linear regression method. We compute both importance (Gradient Boosting Regression with arboreto) and correlation (spearman) scores for all region-gene pairs having a distance between the region and the TSS within the range of 100 to 50k base pairs. In the third step, we extract similar importance and correlation scores as in step two, this time contrasting the gene expression values between the TFs and target genes. For this step, we directly use the output derived from pyscenic analysis with default parameters included in the VSN pipeline (Van de Sande et al., 2020).

Last but not least, the critical step required to finalize the eGRN inference is to prioritize the edges characterized in previous steps that have the highest probability to be functional. First, we dichotomize all edges into activating or repressing functional categories based on a Spearman’s **ρ** threshold of 0.03 (activating if **ρ**>0.03, repressing if **ρ**<0.03, other edges are discarded). Similar to the method from (Aibar et al., 2017), we then generate sets of refined region-to-gene edges via multiple pruning methods based on importance scores. The methods used in this analysis are BASC binarization (Hopfensitz et al., 2012), quantiles (top .75, .80, .85, .90 and .95 quantiles for all regions associated with a given gene) and top highest importance (top 5, 10 and 15 regions associated with a given gene). From these sets of regions, we include specific TF information by associating a TF-to-region edge to each region present in one or multiple TF’s cistrome, from both direct and extended categories (see *cistaret* analysis in step one).

The fourth and last step of the GRN inference consists in pruning the resulting list of TF-to-region-to-gene edges based on the TF-to-gene importance ranking obtained from step three. To do so, we retain the leading edges from a Gene Set Enrichment Analysis (GSEA) (Subramanian et al., 2005), taking the TF-to-gene importance from step three as ranking values and the target gene from step one-two as the tested gene set. Lastly, we unified all the leading region-to-gene edges from the same cistromes and functional category into a common eGRN. This way, for each TF, we preserve the distinction between activating, repressing, direct and extended eGRNs.

### eGRN analysis

Having access to eGRN information allows us to further explore our multiome dataset. We can score an eGRN activity at the single-cell level, for both gene expression and chromatin accessibility using the *AUCell* module from the SCENIC pipeline. An eGRN will have a high AUC score in a cell if its set of target genes and regions are overall ranking high in terms of gene expression and chromatin accessibility respectively. To limit the impact of population size, we compute AUC scores on 150 pseudo-bulk cells from each cell type, each generated from a random selection of 15 cells for the same cell type. We define a set of high-quality eGRN by computing Pearson’s correlation between TF expression and eGRN activity across the pseudo-bulk cells and retain the regulons with an absolute **ρ** higher than 0.2. We additionally remove extended eGRN from the selection if the direct eGRN from the same TF is already present. The final list of eGRN comprises 98 TFs and 147 eGRNs, and is available in Supplementary Table 2, along with correlation results.

In addition to pseudo-bulk, we also derived AUC scores from the full multiome dataset at single-cell level. These scores can be queried via the SCope platform (https://scope.aertslab.org/#/WingAtlas/*/welcome) and are shown as an example in Figure 2.b. Using these AUC scores, we used the method presented in (Suo et al., 2018) to compute a Regulon Specificity Score (RSS) for each eGRN. This RSS indicates whether the eGRN target gene and region sets are specifically active in one cell type (Supplementary Figure 3.a). We retrieve the top-scoring eGRN for each cell type to build the Figure 2.c (gene-based RSS) and Figure 3.a (region-based RSS, NES scores retrieved from the cistarget analysis).

### Data availability

In-house sequencing data and aligned matrices have been deposited in GEO (accession code GSE205401). Public raw scRNA sequencing files for wild-type wing discs were accessed from the GEO accession codes GSE155543, GSE133204 and GSE127832. Public raw scRNA sequencing files for wild-type eye-antennal discs were accessed from the GEO accession code GSE141590. Public raw scRNA sequencing files for 14-day induced scrib^-/-^ eye-antennal discs were accessed from the GEO accession code GSE130566. Public raw ATAC sequencing files for wild-type and *ras*^*V12*^ *scrib*^*-/-*^ eye-antennal discs were accessed from the GEO accession code GSE59078. Single-cell gene expression and chromatin accessibility patterns for all analyses are accessible through the UCSC Genome Browser (http://genome.ucsc.edu/cgi-bin/hgTracks?db=dm6&hubUrl=https://ucsctracks.aertslab.org/papers/wing_disc_wounding/hub.txt) and through the SCope platform (https://scope.aertslab.org/#/WingAtlas/*/welcome), along with eGRNs and their associated activity scores and target features. All other data generated are included in supplementary files.

## Supplementary Figures

**Supplementary Figure 1:**
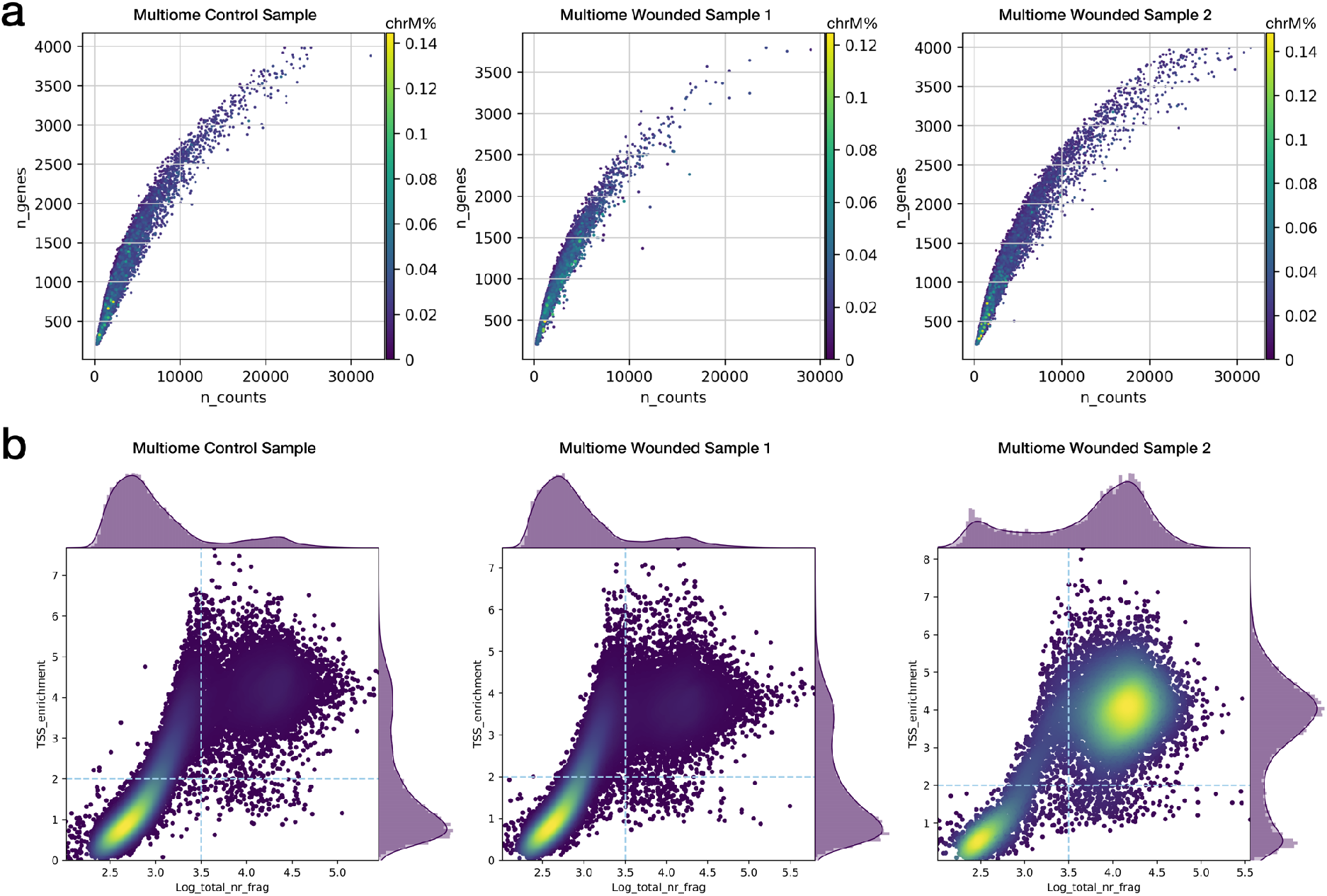
**(a)** Distribution of the number of detected genes and total read counts for the multiome scRNA-seq assays. Color represents the proportion of mitochondrial reads per cell. **(b)** Heatmap distribution of TSS enrichment score and total fragment counts for the multiome scATAC-seq assays.

**Supplementary Figure 2:**
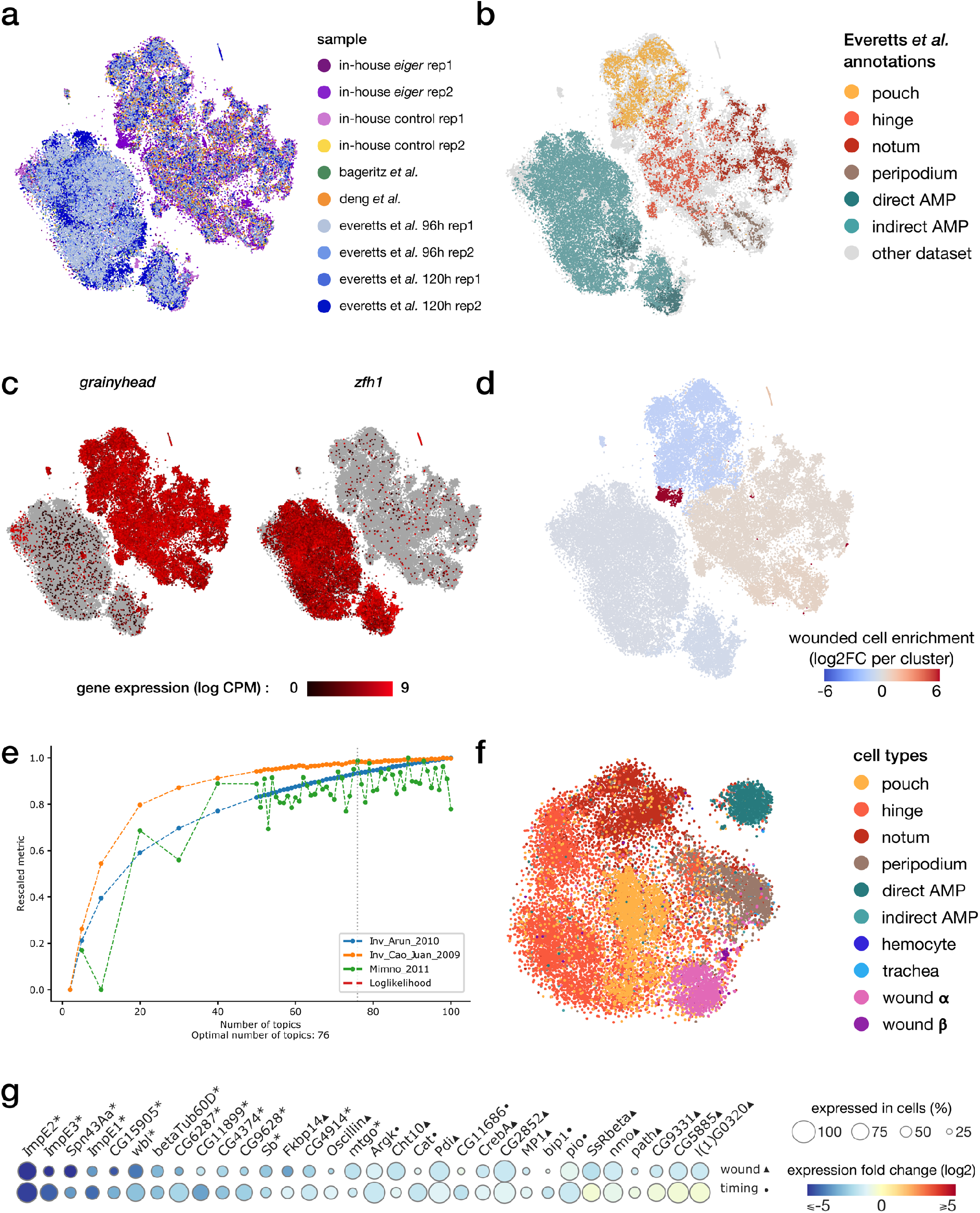
**(a-b)** Harmony-corrected tSNE visualization of the scRNA wing disc atlas, colored by the sample of origin (a) and annotations from Everetts *et al*. (b). **(c)** Gene expression patterns of the epithelial marker *grainyhead* and the myoblast marker *zfh1*. **(d)** Enrichment of cells originating from the wounded samples for each cluster. We note an imbalance in wild-type/wounded cell proportion in the pouch cluster (wound-depleted) and the wound cluster (wound-enriched). **(e)** cisTopic selection metrics across all tested LDA models. The selected model with best coherence scores comprises 76 topics. **(f)** tSNE representation of the topic embedding generated from the multiome scATAC data, colored by cell type identified from the scRNA view. **(g)** Relative expression (log2FC) of markers of wound response (1st row, wounded/wild-type, ▴) and/or developmental timing (2nd row, 96h/120h after egg laying, •) in the notum domain. Shared markers are marked by an asterisk.

**Supplementary Figure 3:**
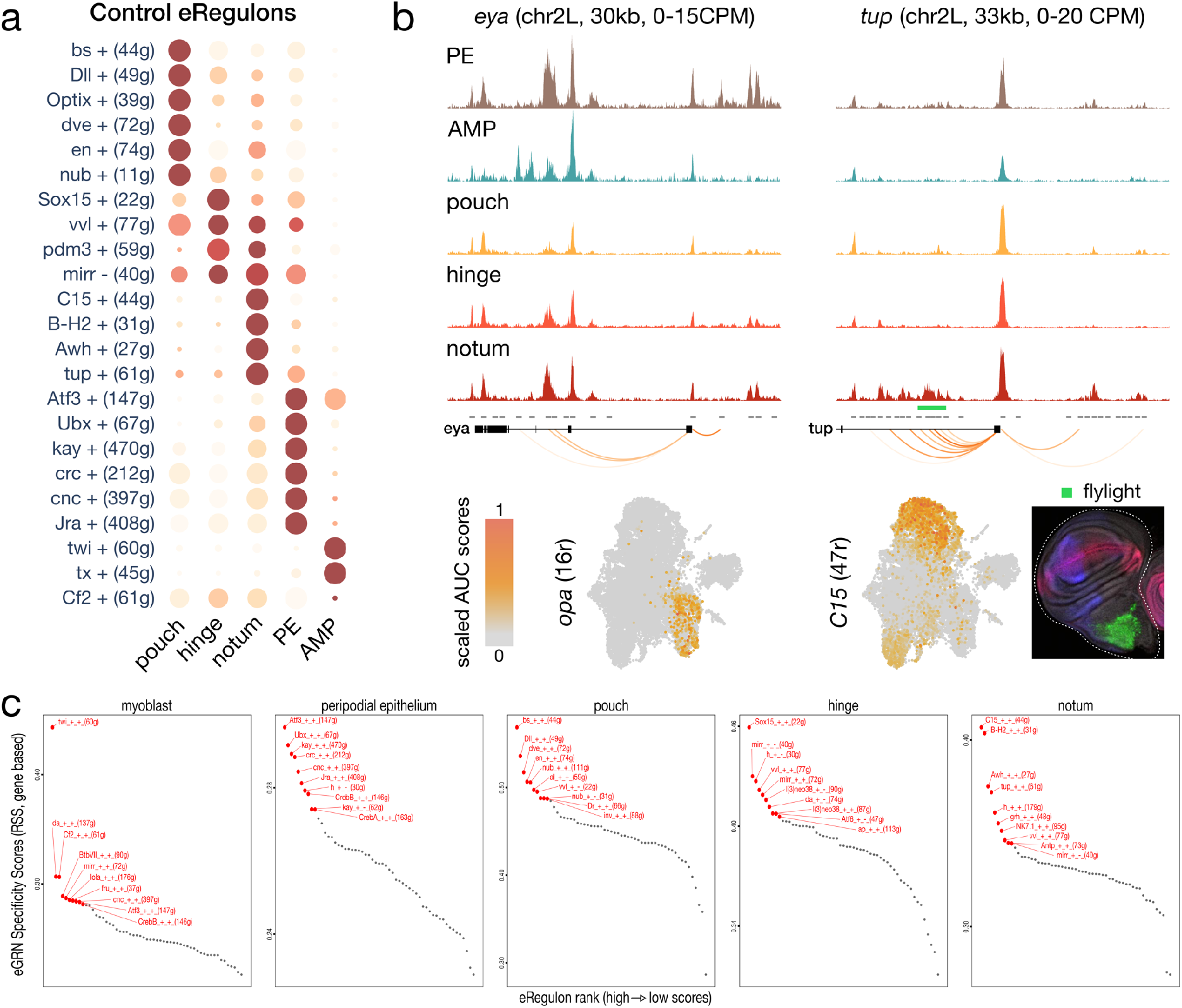
**(a)** Dotplot of the average gene-based activity of selected eGRNs with highest cell type specificity, for wild-type conditions. **(b)** Aggregated wild-type chromatin accessibility signals around a target gene of the *opa* (left) and *C15* (right) eGRNs. The target regions are significantly correlated with expression of each TF (orange arcs), and *C15* targets overlap a flylight reporter (green rectangle) expressed in the hinge. The eGRN activities (UMAP) are localized in the peripodial epithelium (*opa*) and the hinge (C15). **(c)** Regulon Specificity Score of selected eGRNs for each cell type, based on wild-type gene expression. Top 10 eGRNs are shown in red.

**Supplementary Figure 4:**
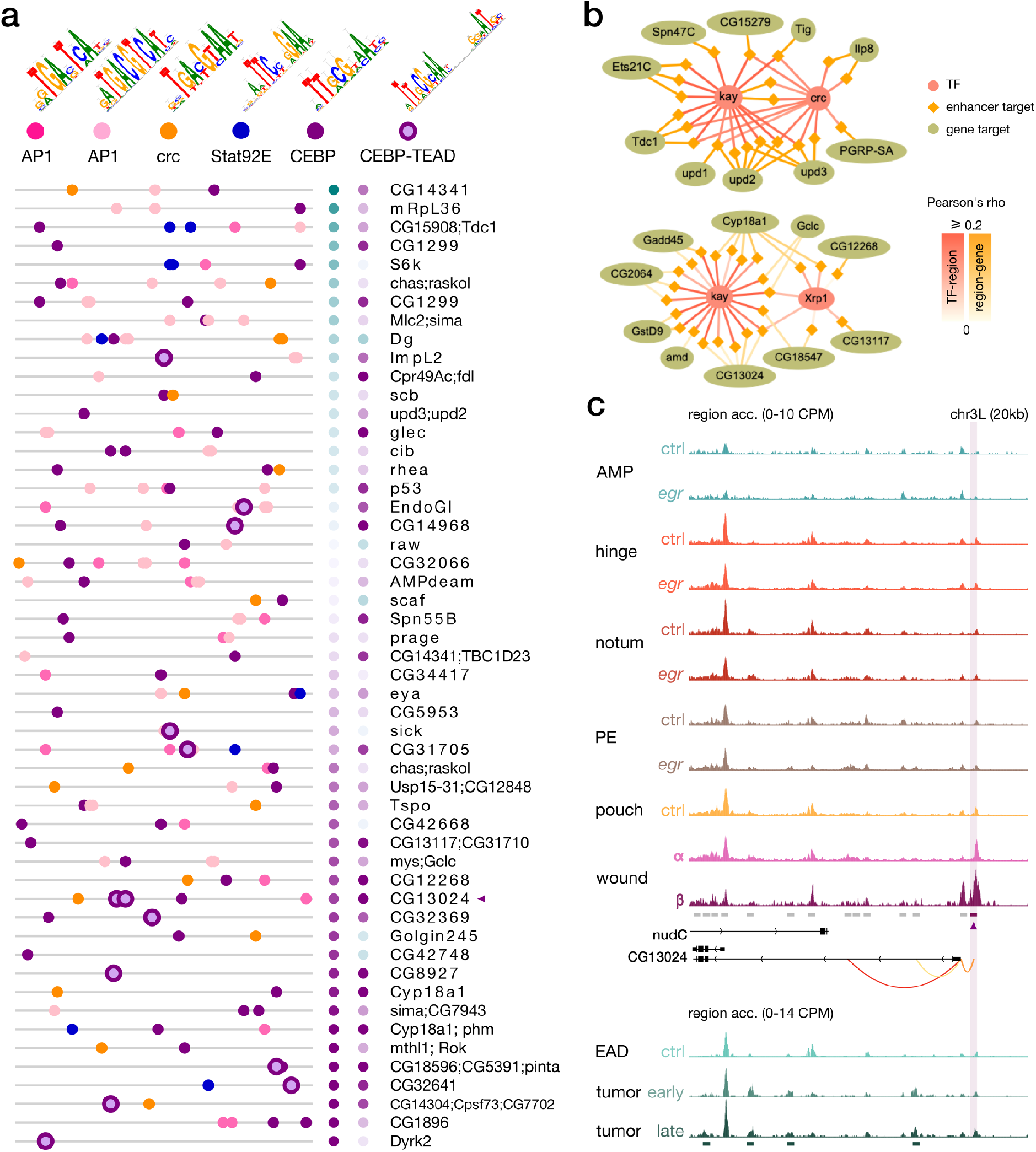
**(a)** Feature maps of six types of TF motifs on enhancers targeted by the C/EBP-like eGRNS. AP-1 bindings are homogeneously present, while C/EBP-TEAD motifs are enriched in targets with higher activity in **β** compared to **α**. **(b)** Example targets regions (lozenges) and genes (ellipses) of eGRNs active in the wound populations **α** (left, *kay* and *crc*) and **β** (right, *kay* and the *Irbp18* partner *Xrp1*). Represented targets are the highest-scoring markers of each wound population. **(c)** Aggregated wild-type, wounded and ras^V12^scrib^-/-^ chromatin accessibility signals around a target enhancer and a target gene (CG13024) of the C/EBP eGRNs. This enhancer comprises a C/EBP-TEAD dimer motif hit as shown in the first panel (purple triangle). Orange arcs represent region-to-gene edges and green rectangles represent the signature regions of the ras^V12^scrib^-/-^ tumor model.

**Supplementary Figure 5:**
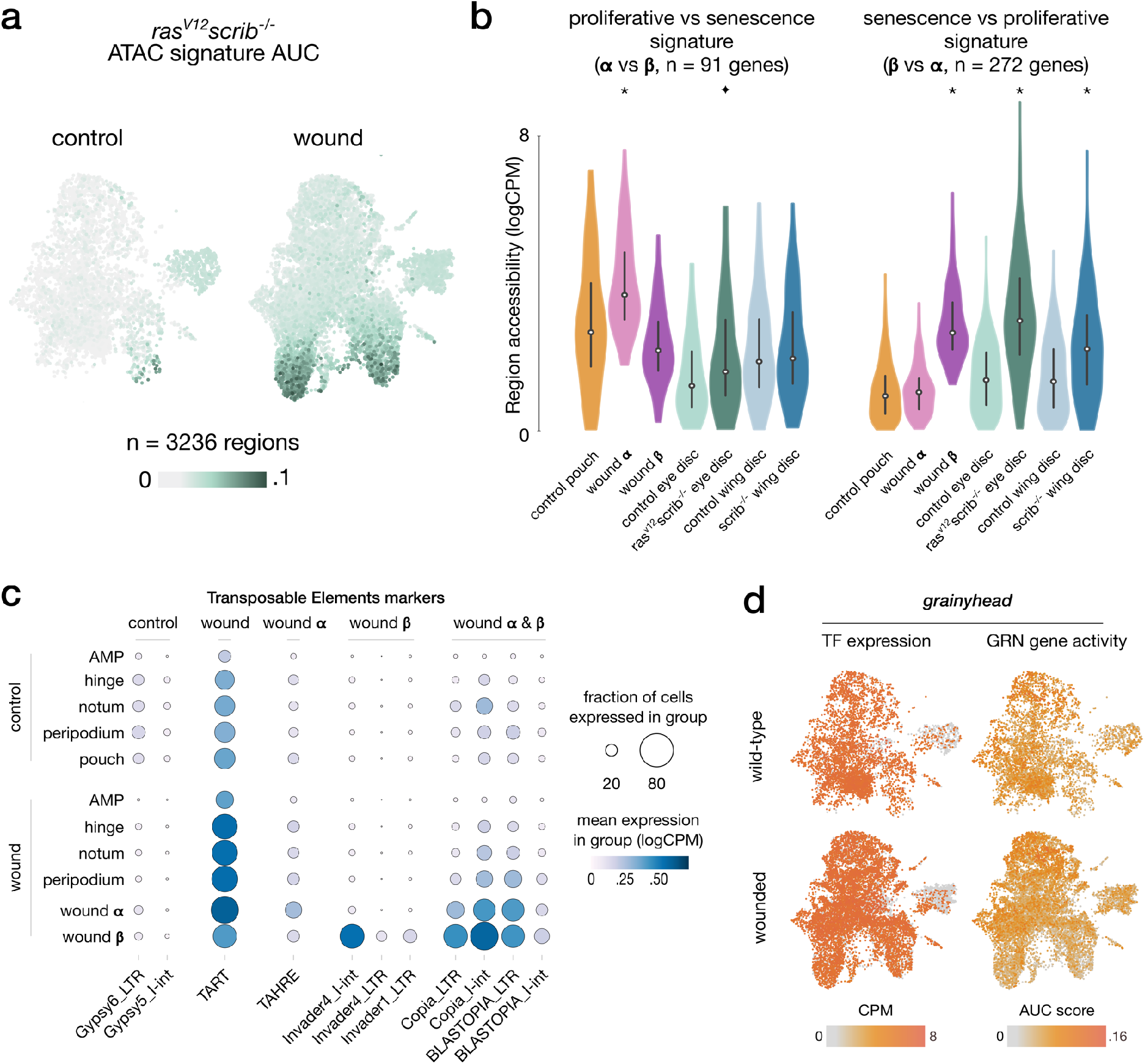
**(a)** Scaled activity of the *ras*^*V12*^*scrib*^*-/-*^ signature inferred from bulk ATAC data, scored using AUCell on wing disc ATAC data. **(b)** Distribution of gene expression measures of the gene signatures from wound **α** and **β** contrasted against each other across aggregated signals, asterisks (p-adj<10e^-3^) and star (p-adj<5*10e^-3^) denote significant enrichment compared to wild-type (one-tailed t-test). We note a higher accessibility of wound **β** signature in the *ras*^*V12*^*scrib*^*-/-*^ ATAC signal. We also find a high accessibility of wound **α** signature in wild-type pouch ATAC signals, indicative of pouch specification markers present in this signature. **(c)** Results of the differential analysis of scRNA signal in Transposable Element regions (p-adj<10e^-3^, log2FC>1). We do not find a strong global change in TE signal between wild-type and wounded conditions. However, we note a clear increase in scRNA signal in the population **β** for the Invader4_I-int retrotransposon. **(d)** Activity of the grainyhead TF and its eGRN-associated target genes in wounded and wild-type conditions.

