## supplementary_tables for "Shared enhancer gene regulatory networks between wound and oncogenic programs": readme.txt

sup_table_1 : markers of the atlas cluster (leiden_resolution = 0.5), using scanpy rank_genes_groups, p_adj cutoff 0.01, log2FC cutoff of log2(3)
sup_table_2 : eRegulon metadata, orthology > 0.6, no min target set size, keep ext. if no direct, 0.2 minimum rho
sup_table_3 : markers of the wound populations (alpha vs. beta and both versus pouch), using scanpy rank_genes_groups, p_adj cutoff 0.01, log2FC cutoff of log2(2)
sup_table_4 : regions and TF motif hit information used to construct featuremaps, incluse the CEBP-specific target genes
